# Differential Expression Analysis of Cannabis sativa response to thermotherapy of Hops Latent Viroid (HLVd) infection and clearance of the viroid in Tissue Culture Micropropagation

**DOI:** 10.1101/2024.04.06.588422

**Authors:** Anthony Torres, Chris Pauli, Carolina Sarmiento, Chris Zalewski, Reggie Gaudino

## Abstract

Increased farming and cultivation of *Cannabis sativa* is rapidly pushing Cannabis (*Cannabis sativa* L.) towards becoming a commercial agricultural commodity. Large-scale cultivation facilities maintain thousands of clonal varieties of recreational and medicinal cannabis and there is a strong market-driven motivation to maintain a commercial pipeline of clean healthy vigorously growing plants free of pathogens. However, mass production and high-capacity cultivation create an environment that is susceptible to highly transmissible pathogens and infectious entities such as Hop Latent Viroid (HLVd). From nurseries to cultivation facilities, it’s become increasingly important to maintain a routine testing regimen and ensure cultivation facilities are HLVd-free environments. One method to address the problem of infected plants is to perform thermotherapy on *C. sativa* explants under tissue culture conditions and isolate clean tissue to multiply productive and healthy mature plants. We carried out a novel thermotherapy method using tissue culture in 5 varieties of type III cannabis that were HLVd positive to document the degree of success of the treatment at the RNA level. We observed that following thermotherapy treatment we were able to decrease the level of HLVd positive tests in select varieties and observed some varieties were highly susceptible and unable to clear the viroid. Plants were tested using a one-step RT-qPCR method, developed and validated, in part, along with this work, and present the results as well as an exploratory transcriptome analysis of an internally developed variety, AnnaLee, which tested negative for HLVd following thermotherapy treatment, and explore possible genes of interest for viroid infection, clearance, and mitigation.

**Author Summary:** RT-qPCR and transcriptome analysis of Hops latent viroid (HLVd) in infected and non-infected *Cannabis* varieties. A thermotherapy procedure was conducted on HLVd infected *Cannabis sativa* meristem tissue maintained through tissue culture micropropagation techniques. Total RNA was isolated from the cultured plantlet stocks and evaluated by a real-time reverse transcriptase assay for HLVd. Infection status post thermotherapy was assessed, and viroid-free plants were maintained and subsequently tested. A single thermotherapy-treated cultivar, Anna Lee was selected for transcriptomics, and an analysis of the genes that were differentially regulated in infected and non-infected treated plants is discussed.

## Introduction

Hops Latent Viroid (HLVd) is a naturally occurring, 256 base pair circular and functional RNA oligonucleotide that has been characterized as a viroid with inherent structure, function, and unique pathology (1, 2), (3). It was first identified in the late 1980s in hop fields and has been reported to have a remarkable function and capacity to spread across large hectarage in a rapid progression, causing “dudding” during flower development, decreased yield, and adverse effect on cannabinoid and terpene production (1), (4), (5). Incidentally, there is a notably high degree of sequence similarity between the sequences of the viroid available from Hops and *Cannabis sativa* (1),(3), (6),(7).

The pathology of HLVd in *C. sativa* has been reported (3) and there is an increased understanding that the symptoms associated with HLVd are unique, elusive, and then drastically noticeable upon environmental conditions or inductive stresses, like flowering development, that begin a symptomatic cascade (7), (8). HLVd can be vector driven or mechanically transmitted during vegetative processing practices (6). The symptoms associated with this reported viroid are observed in the stem, leaf and flower structure (7). Markedly, the production output is also noted to decrease yield and secondary metabolite production, which often are the plant’s major line of defense against pests/predators and primary commercial value (6),(9).

In recent years, HLVd reporting, and observation has led to increased awareness of the potential for HLVd to cause devastating consequences in *C. sativa* by affecting growth patterns and subsequent harvest/production potential (3),(6),(7) demonstrating the significant agricultural and economic implications of unchecked viroid spread. HLVd can be elusive in its incidence and progression. Some say it’s best found in petioles or meristematic tissue (10), recent evidence suggests roots may be the best tissue to sample for assay consistency, but these studies are still ongoing. Monitoring and surveillance methods were internally developed to manage HLVd infection in our own populations and described in this study. Reverse transcriptase or Loop mediated isothermal amplification-based approaches have shown, with great success, the ability to accurately and sensitively detect the viroid. (8), (11), (12). Despite the viroid maintaining a considerable secondary structure (1), well-designed targeted assays measuring stable amplicon products of the viroid provided a robust approach for identifying this relatively small viroid given that an amplicon will query most of the viroid. Pokorn previously identified the upregulation pattern of a lipooxygenase stress marker in HLVd positive hop plants (13) and Punja demonstrated the threat the viroid poses in Cannabis cultivation, categorizing symptomology and effect (6). We confirm some of those findings in this study from a molecular perspective and have found that expression targets of interest have been implicated, however the exact mechanism underlying HLVd’s transition from asymptomatic to symptomatic remains unknown.

We sought to better understand HLVd in the context of micropropagation and test whether there is an ability to clean it from infected plants using thermotherapy treatment (14) and further identify which genes, if any, were potentially differentially regulated upon HLVd infection in meristematic tissue. Accurate assays are essential for early detection of the viroid and more understanding is needed about the genes that are potentially affected in a HLVd infected Cannabis plant. We tested over a dozen varieties and determined that there is a potential for viroid clearance using thermotherapy in micropropagated plants and follow up monitoring/testing. There were varying degrees of success depending on the variety. We also observed the treatment was reproducible and long lasting several months following treatment. In one variety, we carried out Nanopore cDNA sequencing on replicates of negative and positive samples and identified a range of potential gene targets that fit the profile of reported symptoms and potential genes that would be involved in structure and integrity of tissue infected with HLVd. We have made key observations that may shed light on the potential points the viroid may be hijacking biochemical pathways, hinting at its pathogenetic strategy. These results contribute to the understanding of HLVd, its molecular pathology, and methods that can be employed to mitigate a major pathological issue in the field.

## Results

Thermotherapy treatment and Outcome of HLVd clearance following thermotherapy We sought to deploy the HLVd RT-qPCR detection assay in an analysis of HLVd surveillance and tracking in a tissue culture environment of meristematic tissue regenerants. We observed presence of HLVd in the tissue culture population from plants coming in from external uncontrolled environments (3^rd^ party testing data from the period between May 2020 through October 2020). Plants with a confirmed infection of Hop Latent Viroid (HLVd) are put through thermotherapy and meristem isolation, post treatment, the meristem-derived plants are tested twice to confirm a negative result of HLVd. The program started in June 2020, and as of January 2021, six varieties have been cleaned and multiplied into combined amount of 1,754 plants of stock.

The plants appeared to tolerate the thermotherapy conditions well with survival rates greater than 70% after heat treatment. The treatment was expected to eliminate the viroid by regenerative clearance relying upon cell division occurring more rapidly under thermotherapy conditions with inhibition of viroid replication due to temperature treatment resulting in certain subpopulations of the newly divided and cleaned meristem containing no viroid. We performed an initial test following treatment on a randomized 8-28% sampling of the surviving meristem plantlets (Table 2). We observed a distribution of regenerative clearance in survivors that allowed a calculation of the number of clean plants (HLVd negative) by the number of total tested plants to determine the percent cleaned by the thermotherapy method. We observed some varieties were able to show significant regenerative clearance and improvement. Athena and Valarie showed a high degree of viroid clearance at 94% and 100% clearance respectively. AnnaLee, FRB1.4 and Wife showed similar degrees of viroid clearance and an intermediate level of post thermotherapy viroid eradication, with 62% viroid clearance for AnnaLee and 69% for FRB1.4 and 50% for Wife. Hybrid 9, Hybrid 5, EarlyPearly showed a moderately low level of viroid clearance with 26%, 14% and 14%, respectively, and two varieties showed little to no clearance of the viroid, which were Abigail with 3% clearance and S1-4 with 2% clearance. Two varieties, RiverRock and PureCBG showed no clearance following thermotherapy and were removed from the study. (Fig 2 A.) A second test was performed on the negative testing plants 10 weeks after the initial test and the viroid cleared plants remained cleaned in their environment (Fig 2 B.) Results were reported by cycle threshold (Ct) detection of the HLVd probe complementary target and internal reference gene 26S (Fig 1). Fig 1 shows a box-plot analysis of the Ct results measured in the RT-qPCR assay carried out for initial testing (1 A) and follow up testing (1 B). HLVd negative testing samples observed no Ct and are noted as a line at the zero baseline of the plot (Fig 1 A). HLVd positive plants (Fig 1A Green) had a median result that varied by variety but in general viroid detection was observed between 18-32 cycles. Reference gene measurements were given again by zero Ct for HLVd probe (marked as line at zero baseline for boxplot) and relatively uniform Ct of 26S between varieties and were between 18-22 cycles in both initial testing (Fig 1 A) and follow up testing (Fig 1 B) of the thermotherapy treated plants. Mean Ct values for HLVd positive cultivars were between 19-32 cycles, and mean ct values for the corresponding reference gene was between 14-22 cycles (Table S.2.T qPCR Statistics). Delta Ct was ranged –1.137-16.589 across cultivars. One-way ANOVA analysis of the difference in mean Ct values measured between positive cultivars revealed there was a statistically significant difference in HLVd infection as given by the mean Ct measured between cultivars with 11 degrees of freedom, F value 127.8, p<0.001. The varying degrees of HLVd clearance demonstrate an underlying complex mechanism of clearance as well as the viroid’s mechanism of action. We followed up with choosing an intermediate candidate of viroid clearance for by choosing the variety AnnaLee, a candidate that showed intermediate viroid clearance, for follow up sequencing and differential gene expression analysis using cDNA libraries isolated from HLVd positive and negative tissue.

**Figure 1.**
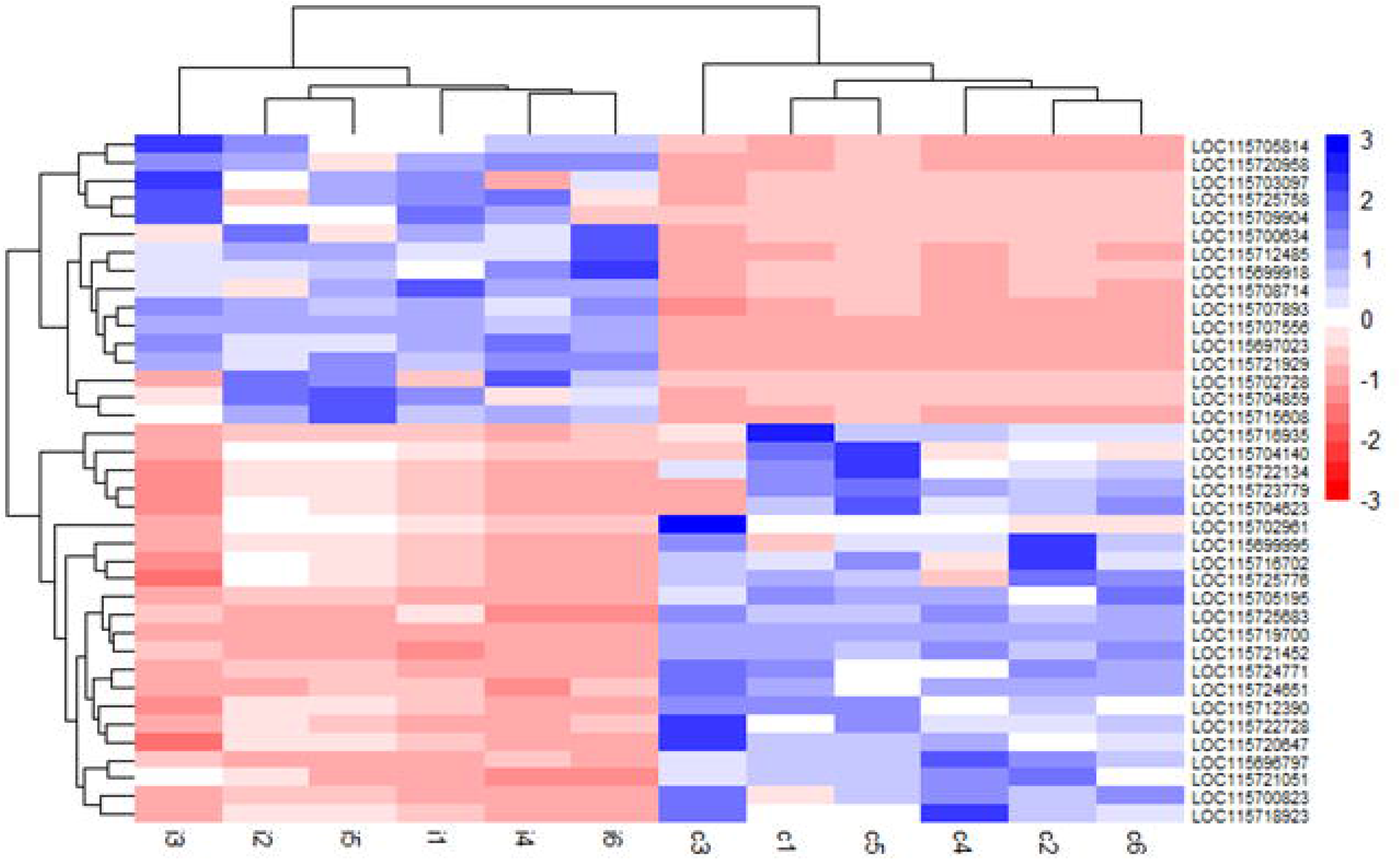
Ct Detection of HPLVD Clearance following initial thermotherapy (top) and post thermotherapy and testing selection (bottom). Blue: internal positive control targeting 26S rRNA internal reference gene.Green: Plants that were unsuccessfully cleared of the viroid. Line at Zero: Plants that had been successfully cleaned had no detectable amount of HLVd. Error bars and median calculated based on distribution of Ct values in detections of HLVd positive and Reference Positive signals. A. Initial Screening of Abagail (n=33), Amber (n=54), AnnaLee (n=39), Athena (n=16), EarlyPearly (n=42), FRB1.4 (n=42), Hybrid5 (n=42), Hybrid9 (n=38), PureCBG (n=23), RiverRock (n=21), S1.4 (n=42), Valarie (n=23), and Wife (n=12). B. Results of confirmatory screening following selection of Abigail (n=1), Amber (n=15), AnnaLee (n=28), Athena (n=12), Hybrid9 (n=10), and Wife (n=6). (Raw Ct Data in Supplemental Table S.1.T)

**Figure 2.**
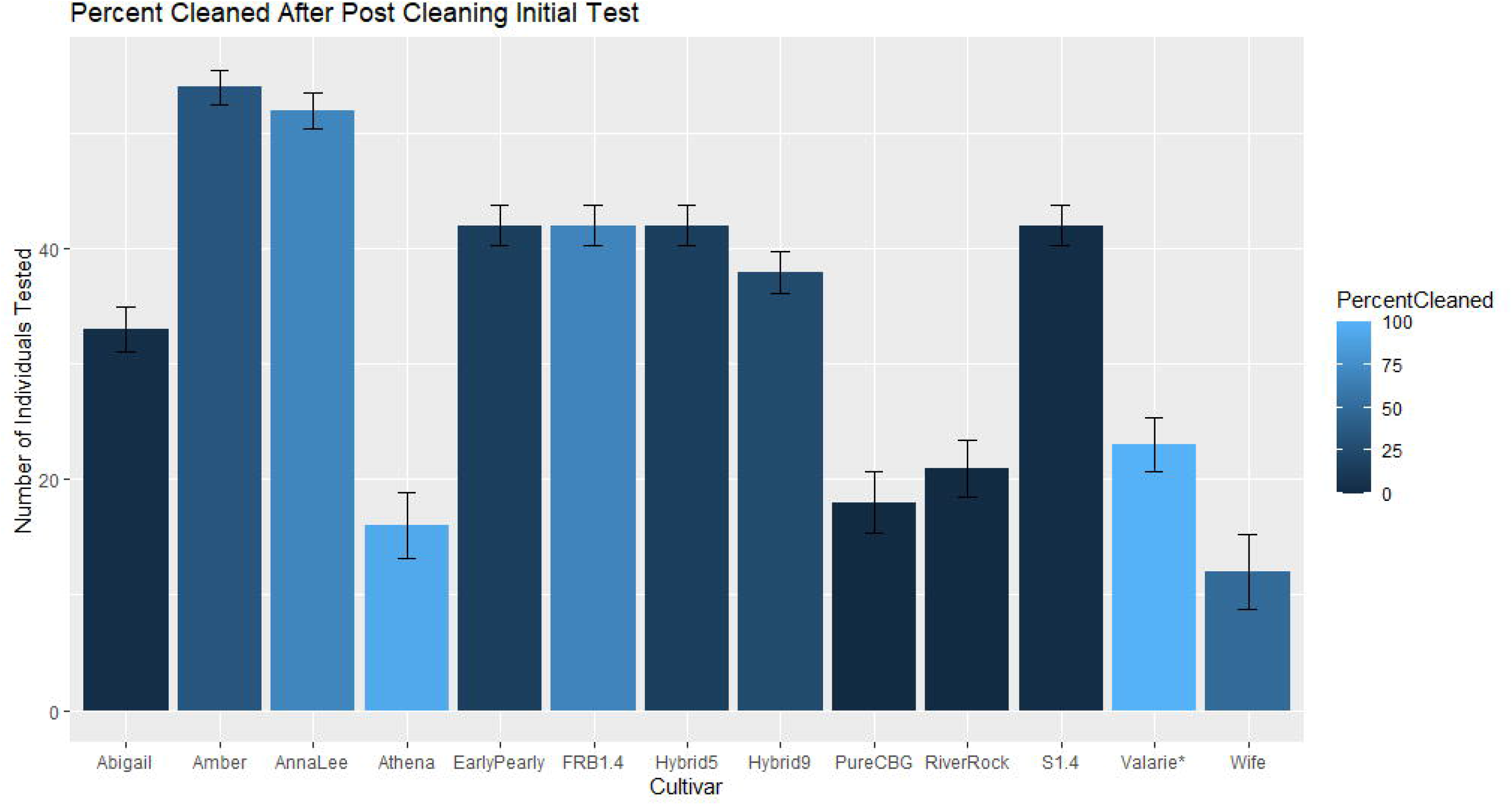
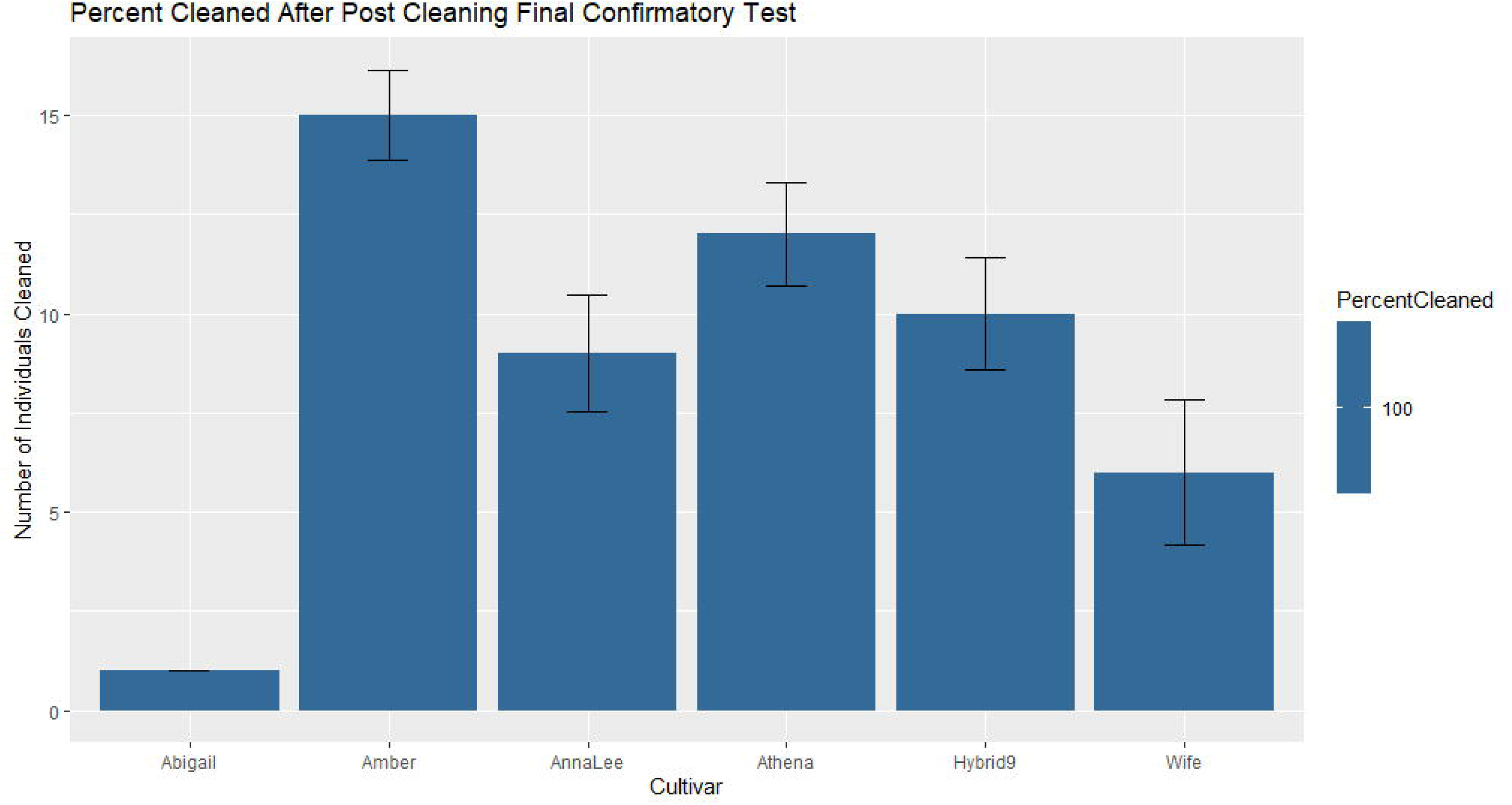
A. Number of plants that had been cleaned and the percentage cleaned of HLVd after initial testing post thermotherapy separated by variety. B. Number of plants that remained 100% clean and HLVd negative in subsequent testing three months post thermotherapy following initial testing. Error bars calculated by the standard deviation and standard error of number of negative individuals in the population test set.

### HLVd detection and recovery by sequencing

The raw long read cDNA nanopore reads were basecalled and mapped to a reference (Accession # GCF_900626175.2) (15). HLVd full length viroid was detected in the positive nanopore cDNA libraries and was absent in the negative cDNA libraries. As previously described by Pokorn et al., three potential gene targets of HLVd were found in *Humulus lupus* (13). From the full-length transcripts pulled down from Nanopore cDNA sequencing, we identified full length transcripts of several family members of the LOX gene family and identified one to have a notable differential expression profile in the HLVd tissue vs non infected. Other LOX genes showed baseline expression levels. It is possible that the one LOX family member identified was a homolog of the *H. lupus* candidate gene that Pokorn reported (Pokorn 2017). LOX was not of the most highly expressed genes in this tissue profile, however, the difference in expression patterns was measurable. We also identified the 256nt HLVd sequence despite its relatively short sequence. The full dataset of transcripts produced by Nanopore cDNA sequencing of *C. sativa* meristematic tissue used in this experiment are published here (Bio project number PRJNA1083500 and SRA reads include 6 replicate clean Anna Lee tissue libraries (SAMN40250909-SAMN40250914) and 6 replicate infected Anna Lee tissue libraries (SAMN40250915-SAMN40250920)):

### Differential Gene Expression (DGE) analysis

The replicated gene expression libraries of both the control (non-infected) and HLVd-infected were analyzed using DGE to identify a total of 74 genes that were differentially expressed (Supplemental Table S.3.T). Of those 74 genes, 55 of them could be assigned gene annotations using the Cs10 RefSeq (Accession # GCF_900626175.2), and 19 of these genes were labeled as “uncharacterized proteins”. Of the assigned differentially expressed genes, 38 genes were considered significantly up-regulated, and 36 were downregulated in response to being infected with HLVd with respect to verified uninfected controls (Fig 3). Due to the controlled design of this experiment, the magnitude of the differences between infected and noninfected libraries can be attributed to the presence of HLVd. These observations were only possible due to the strict temperature, light, humidity, and nutrient control that is part of the tissue cultivation protocol used. We observed upregulation of Expansin genes and cell wall modifiers, along with other genes related to signaling and defense cascades (Table 3-A), as well as downregulation of structural component genes and regulators of primary and secondary metabolism (Table 3-B). We identified a hallmark expression pattern of upregulated genes involved in plant architecture, cell defense, and stress responses upon HLVd infection. Stress response related genes such as cytochrome P450 94A1-like (LOC115722134) and protein SULFUR DEFICIENCY-INDUCED 1 (LOC115712390) showed a nearly 5-fold increased expression in infected tissue (Heatmap i1-i6).

**Figure 3.**
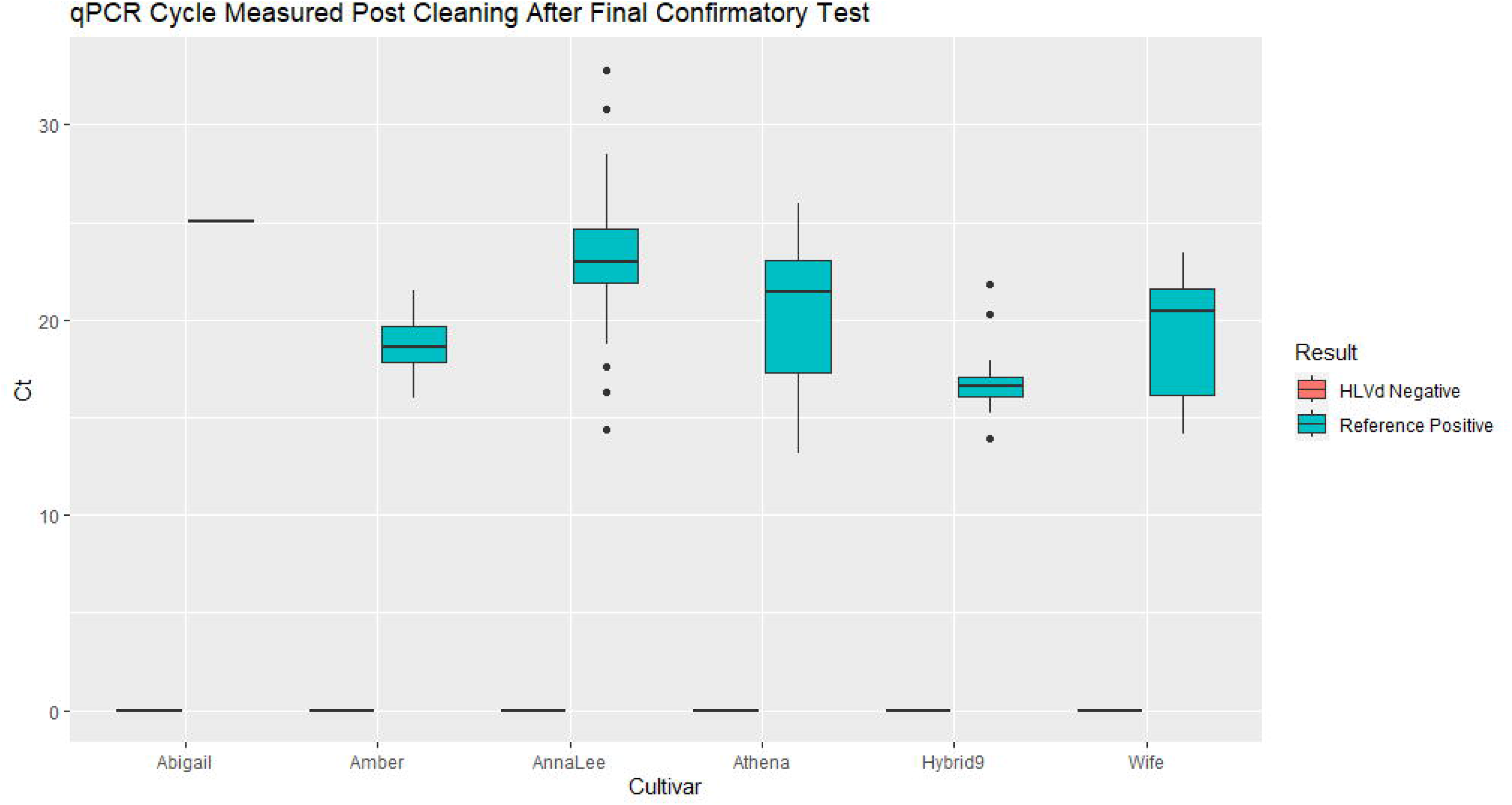
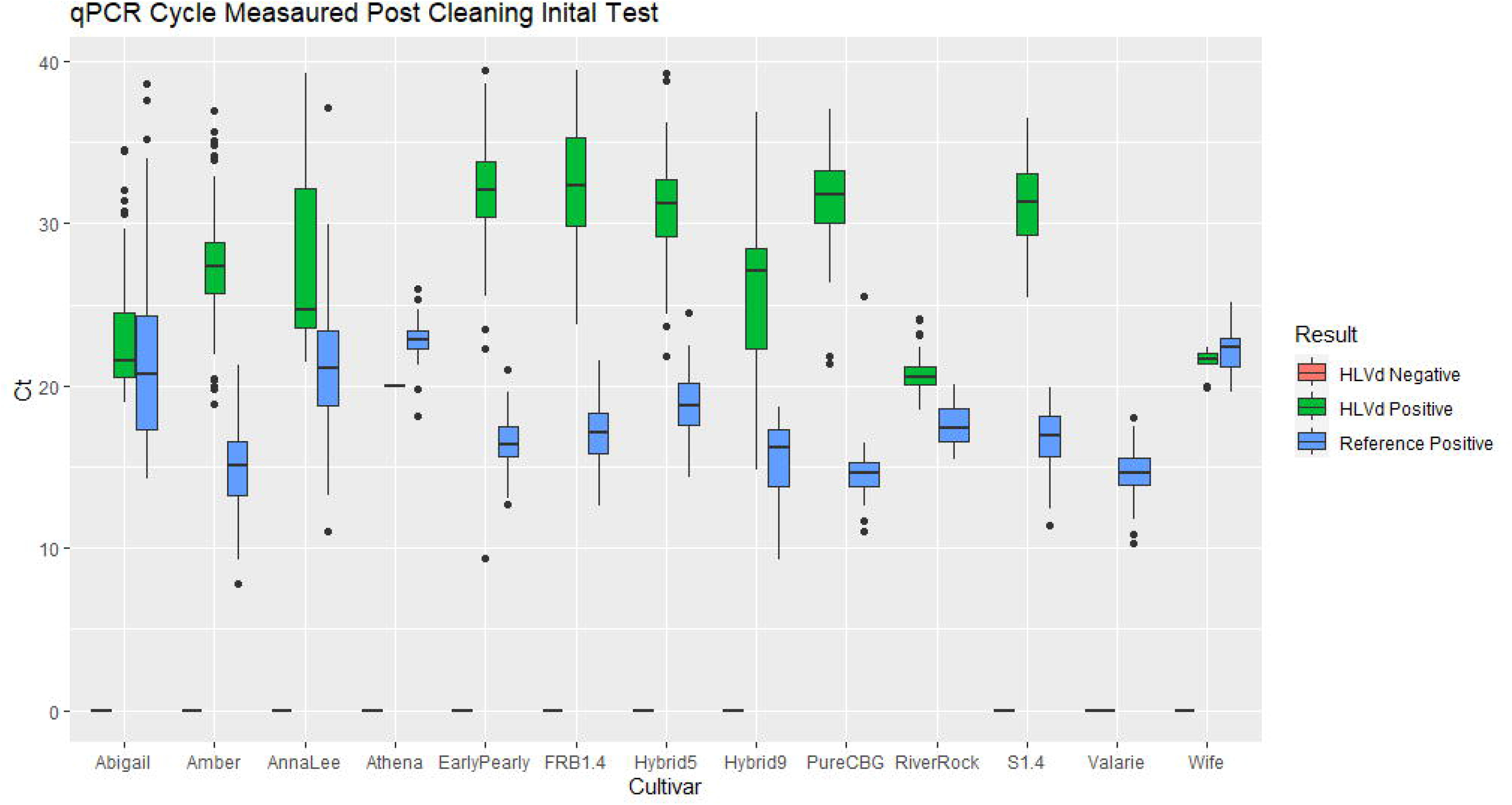
Heatmap of the differential gene analysis from 6 replicate HLVd infected (i1-i6) and non infected (c1-c6) Cannabis meristematic tissue sample libraries. Results visualized as Log Fold change of normalized full length nanopore cDNA read counts in infected vs non infected is represented in color (blue is upregulation and red is down regulation).

Additionally, we observed genes involved in defense and cell surface signaling such as calcium-binding protein CML37 (LOC115718923), probable protein kinase At2g41970 (LOC115699995), ethylene-responsive transcription factor 5 (LOC115717279), and probable xyloglucan endotransglucosylase/hydrolase protein 23 (LOC115724651)that all showed greater than 3.5 fold expression difference, and architecture gene expansin-like B1 (LOC115716935) had nearly a 6 fold increase in expression (Heatmap i1-i6). Conversely, structural integrity genes such as glycine-rich cell wall structural protein 2 (LOC115695646) and genes involved in macromolecule and energy synthesis such as putative lipid-transfer protein DIR1 (LOC115697023), granule-bound starch synthase 1, chloroplastic/amyloplastic (LOC115715608), and ferredoxin--NADP reductase, root isozyme, chloroplastic-like (LOC115725758) showed a greater than a 3.5-fold expression decrease (Heatmap i1-i6). Notably, pathogen defense response genes S-norcoclaurine synthase (LOC115704859) and pathogenesis-related protein 1 (LOC115705814) showed a greater than 4-fold downregulation in the infected libraries (Heatmap i1-i6). All differences discussed were observed in replicate libraries and had p values less than 0.001 (Supplemental Table p values). The results of this experiment highlight the dynamic nature of stress response pathways and expression changes occurring in various plant gene families including notable changes in cell-wall membrane proteins. The largest category of upregulated genes was classified as signaling and defense genes with 5 genes, as well as 4 genes related to stress and detoxification which were also upregulated as shown on TABLE 3 A and B. Additionally, there was a more distributed classification of downregulated genes, with genes related to cell wall structure, lipid metabolism, protein regulation, photosynthesis, and most interestingly stress response and defense proteins.

## Discussion

HLVd was discovered in 1987 in *H. lupus* plants (Accession # NC_003611.1) (5). Dark Heart Nursery demonstrated (3) at least one genotype of HLVd that also infects *C. sativa* through sequencing confirmation (3) (MK876285 and MK876286). Since it’s identification, various studies and assay design strategies have attempted to identify HLVd’s prevalence and genetic variation within cultivation facilities across North America (6), (11), (13). These reports have indicated that over 40% of dried Cannabis flower cultivated in legal Cannabis markets is infected with HLVd, thus a need for an accurate and cheap HLVd genotyping assay is necessary.

The mechanistic action of how viroids display in a plant’s phenotype is slowly being understood (16), and other studies to understand viral load in various tissues and time points within the plant’s life have been published (17), (18). Due to the asymptomatic phenotypes observed in many plants with early-stage HLVd infection, it is extremely difficult to visually diagnose these early-stage infections, which can lead to the rapid spread of these pathogens throughout a cultivation facility without strict phytosanitary procedures. A preponderance of evidence suggests that mechanical transmission remains the major vector of spread through activities such as trimming and cloning with contaminated trimmers (19).

Eradication of the viroid through thermotherapy procedures as the ones performed in this study have been attempted (1), (20) with various degrees of success likely due to genotypic differences in the plants being cleaned, as well as lowering the viral load to an undetectable level and mistakenly determining a plant viroid-free. In hops, it has been shown that thermomutants are likely created while undergoing heat treatment during tissue culture (21), which increases the risk of a false negative on assays that were not designed with the potentially mutable sites considered. Thus, as part of our work, we developed and validated a highly sensitive and dual interrogation assay, which allowed us to more accurately determine plants recently infected, as well as plants that underwent thermotherapy.

Prior to this study there was no definitive answer as to which genes were involved in the mechanism of action of HLVd in cannabis. The results presented herein suggest that numerous differentially expressed genes may play a role in the presentation of symptoms caused by HLVd. For example, stems have been observed during handling to become more brittle after infection with the viroid (1). In our experiment, we observed cell wall and structure-related proteins being up-regulated, cell wall structural protein being significantly downregulated. This could suggest that the lack of the glycine-rich protein or overexpression of the expansin and hydrolase proteins could be responsible for the brittle-nature of HLVd infected plants (22). Similarly, it has been reported that plants infected with HLVd have also been known to be less vigorous and carry an altered internode length compared to uninfected clones of the same cultivar (9), (23). Here we have observed significant differential expression of macromolecule synthesis and energy related genes being downregulated, which may hint at why growth slows during a high viroid load contaminated state. Furthermore, we also observed downregulation of a pathogenesis-response proteins may suggest that HLVd infection results in a change in plant defense responses, which may suggest a potential mechanism of how the viroid avoids the plant immunity and continues to replicate. Profile changes in the identified genes suggest that these candidates play a role in the infection mechanism, the plant response to the infection, and the downstream gene expression changes that result from this infection.

It’s important to note that the differential expression changes observed could be due to other factors rather than transcriptional gene regulation. As has been shown in previous studies, the single stranded RNA of the viroid is hypothesized to potentially interact with transcripts that have been expressed through complementary binding of the viroid and transcript (13) as a recruiting or functional structured RNA molecule (24), which may cause a lack of translation, or a possible degradation of the transcript due to the double stranded RNA complex formed when the viroid binds mRNA. Additionally, it appears that the plant undergoes a clear stress response post-infection which can lead to other downstream gene expression changes as the plant begins to focus energy on defense proteins and signaling pathways involved in pathogen infection. Finally, the authors are unable to rule out that the differential expression observed in this study is due to an unknown mechanism of pathogen identification and a subsequent transcriptional regulation cascade that causes the changes observed.

There are several limitations of this study including that many genes observed in the analysis are uncharacterized due to the lack of annotation in the Cs10 RefSeq genome annotation. Also, a lack of understanding of the precise role of the identified candidate genes is possible due to the orthologous nature of these genes, which calls for knockdown experiments of these targets in infected and noninfected cultivars. Furthermore, due to financial limitations, our study was limited to transcriptomic analysis of replicates from a single genotype, which limits the extrapolation of this data relative to the larger population germplasm of *Cannabis* analyzed in this study. We were unable to perform potency analysis of secondary metabolites in infected tissue due to their culling from the cultivation environment. Future experiments should include analysis of the differentially expressed genes tracked in real time expression experiments over time in a larger test set of cultivars and conditions using a similarly designed study to further understand the relationship between alleles and the presentation of symptoms in HLVd infected individuals as the HLVd infection progresses. Lastly, using novel qPCR markers for the gene targets identified here, a follow up study would be able to study the patterns found here in a wider range of genotypes to understand the common mechanism of gene expression changes in response to HLVd.

Finally, because not much is known about the relationship between viroid load and symptomatic progression of the disease state caused by HLVD, we felt an assay that detected the viroid at very low loads was important in order to be able to evaluate eradication efforts, as well as potentially be able to identify a level at which the plant goes from being non-symptomatic to symptomatic, understanding that threshold level would likely be variable in different cultivars. Our assay demonstrates sensitivity down to femtogram (fg) levels of viroid detection (Supplemental Fig S.A), as demonstrated in reproducible standard curves. Replicates are a staple in *Cannabis* pathology testing as a rule of practice mainly to ensure multiple samplings of the plant are tested. Some have claimed it localizes into specific parts of the plant as well as multiple time points such as changes in the production cycle, however, there is no conclusive evidence to suggest this. Once the viroid has deployed and is in a high copy or replicative state, it is reproducibly detectable in any tissue/sampling. As such, It’s important to be able to test down to extremely low levels to ensure detection of viroid presence in any tissue.

## Conclusion

We demonstrate differential gene analysis of HLVd in infected versus uninfected *Cannabis* tissue which has elucidated putative targets affected by the viroid. We showed that HLVd can be cleared through regenerative clearance using thermotherapy methods, and there is a difference in thermotherapy response for viroid clearance in different varieties. Lastly, we observe that HLVd appears to affect genes involved in cell wall architecture, cell signaling and defense, and growth-related synthesis.

## Methods and Materials

### Plant Material

13 varieties of *Cannabis sativa L*. were assayed over a 6-month period. The tissue culture explants were maintained under proprietary FRB tissue culture conditions during this time, allowing for micropropagation of the variety. *C. sativa* from 12 type III FRB varieties and one type IV variety: Anna Lee, Abagail, Amber, River Rock, Hybrid 9, Hybrid 5, Wife, Athena, Early Pearly, FRB1.4, S1.4, Valarie, and Pure CBG were cultivated in tissue culture test tubes on agar at room temperature in growth chambers under photoperiod of 20 hours of light and 4 hours of darkness. Meristematic tissue was stored in RNA later and transferred into liquid nitrogen for cryogenic grinding using a mortar and pestle.

### Tissue Culture

Tissue culture initiations were started from healthy and chlorophyll rich petiole, leaf, and stem meristematic material free of insect pests and microbial organisms which was grown on Agar #A037 (Caisson Laboratories, Springfield, UT, USA) prepared at 8g/L with 5.32g/L #DKW D2470 and 0.05g/L L-Cysteine #C204 (Phytotech Laboratories, Lenexa, Kansas) and sucrose 30g/L to initiate shoot development and regeneration (25) described in US Patent # US11432487B2. Sterility was qualitatively measured of initiations by sample culture indexing in autoclaved L&W Nutrient Broth (Millipore, MA, USA): Concentrations of 45.22g/L were prepared as instructed on bottle. Culture incubations were grown at room temperature for 7 days and passed or failed based on visual Turbidy indicative of presence or absence of microbial growth on tissue (26). Multiplication and propagation by shoot cuttings and regenerations were carried out through rooting and regeneration of new plants for further cultivation on Agar prepared 8g/L (with 5.32g/L DKW, 0.05g/L L-Cysteine, 30g/L sucrose, 0.74g/L Magnesium sulfate, and 1.5mL/L IBA). Surveillance clean stock analysis was performed routinely with routine indexing and HLVd testing.

### Viroid Eradication in Cannabis by Thermotherapy and Meristem-tip Culture

As described in US Patent # US11432487B2, Stem tip or base of the viroid infected TC plantlet was grown in a test tube on Agar at room temperature growth chambers under a photoperiod of 20 hours of light and 4 hours of darkness. Upon root regeneration, the temperature in the growth chamber was increased to 28-30C and the plantlets were grown for 4-5 weeks. Following thermotherapy, meristematic isolations were performed (25) and grown on Agar (with DKW and L-Cysteine) plates for shoot initiation/regeneration for 2-3 weeks. Isolated shoots were moved to Agar (with DKW, L-Cysteine) test tubes and grown for an additional 8 weeks, with a media transfer after 4 weeks. Following regeneration, meristematic tissue was stored in RNA later at two independent sampling time points, and RNA isolations were performed on liquid nitrogen pulverized tissue. Isolated RNA was tested for HLVd post viroid thermotherapy treatment and confirmed through sequencing.

### RNA isolation

RNA was isolated and purified using the Qiagen Total RNA easy kit (Qiagen, Germany) and Zymo Quick RNA kit (zymo, Emeryville) following the manufacturer’s instructions. RNA was quantified using the Qubit HS-Sensitivity RNA kit and normalized for 1ng total RNA sample input per reaction (Thermo Fisher, Fremont, CA).

### RT-qPCR HLVd Detection

1ng of normalized RNA was used as input into a multiplex RT-qPCR reaction prepared with B-F HLVd and TC primers with the IDT LNR probes HLVd p4 (FAM), HLVd p2 (Cy5), 26S (SUN) Universal probes for RT-qPcR and iTaq Mastermix (Biorad, Hercules,CA).Primers and probes used in this study are listed in Table 1. Ct values were compared against a standard curve of RNA from 10ng-0.00001ng. Reaction and data acquisition were carried out on a ABI Quantstudio5 and analysis was conducted on the Connect Cloud Data Analysis software (ThermoFisher, Fremont, Ca). The Ct values were called by crossing a threshold value for each fluorescent target. Statistics were calculated using the R package (“pcr”) (27).

**Table 1.**
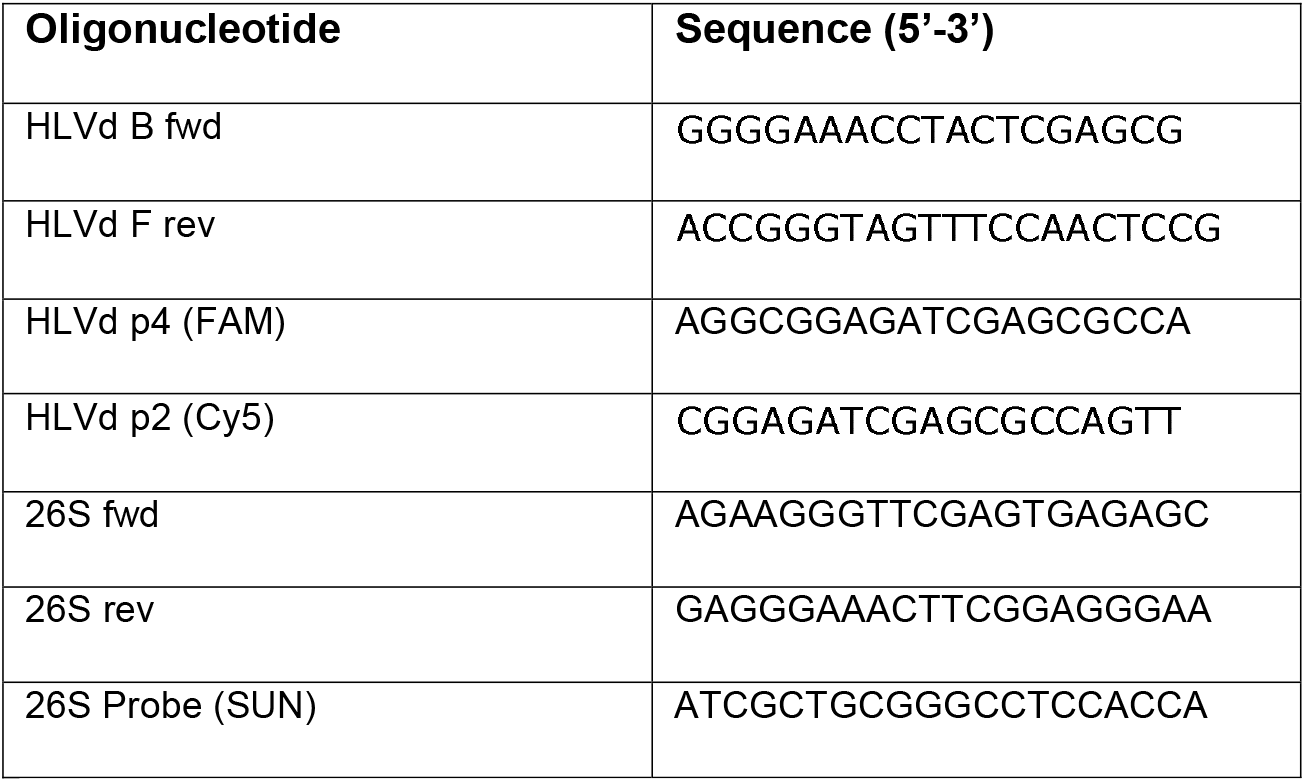
List of Oligonucleotides for dual interrogation testing.

**Table 2.**
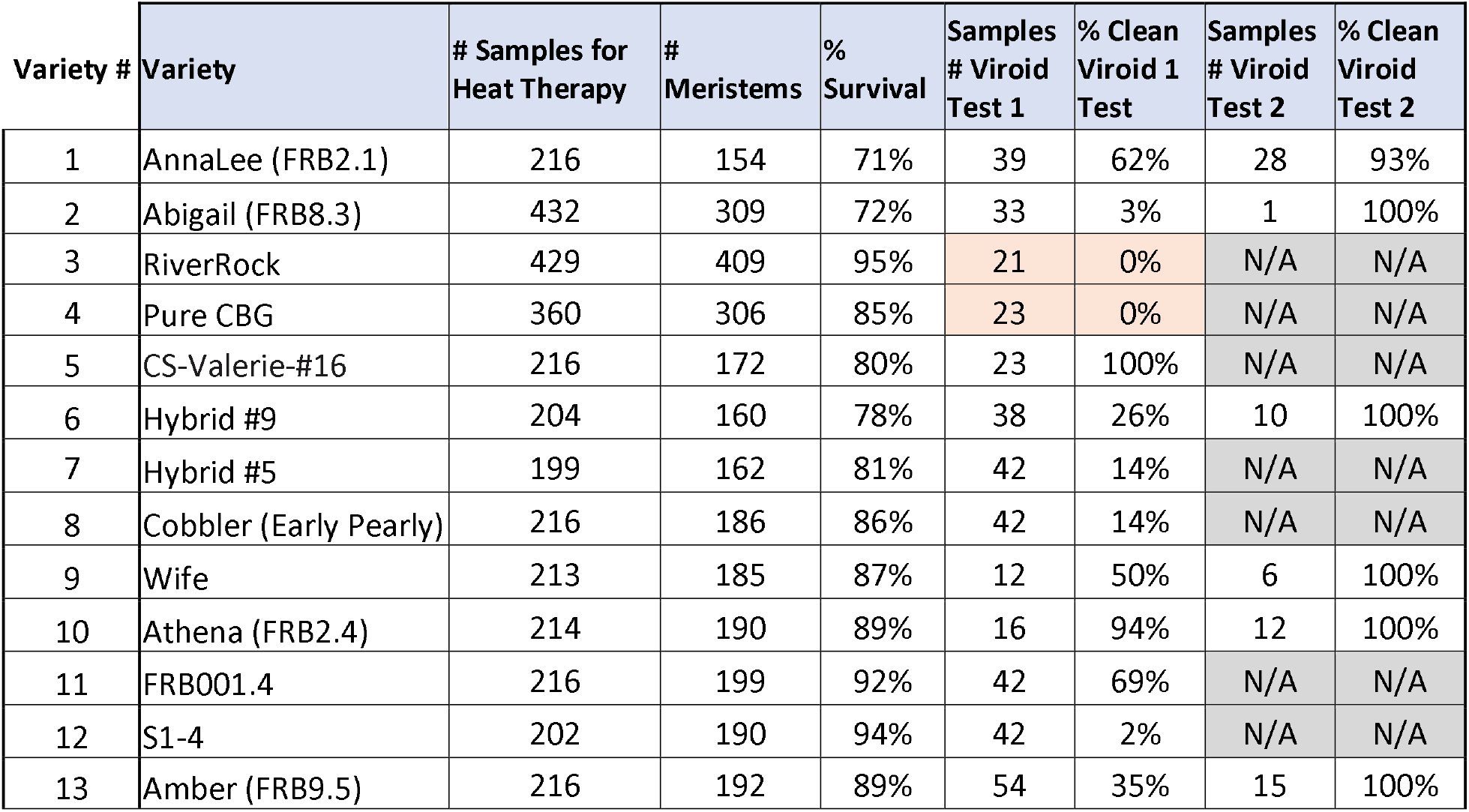
Cultivar Name and Number of varieties tested in this study. We present the number of surviving meristem plantlets post treatment, the number of samples initially tested, and a second round of testing for the negative testing plants. Test 1 and Test 2 were performed 10 weeks (about 2 and a half months) apart and all samples were tested in replicate.

**TABLE 3-A.** Up-regulated Genes in infected HLVd Tissue. Genes significantly upregulated (Greater than 3 log fold change) in infected tissue libraries.

| Activity | Gene ID | Annotated Name |
| --- | --- | --- |
| Cell Wall & Expansion | LOC115716935 | Expansin-like B1 |
| Cell Wall & Expansion | LOC115724651,<br>LOC115712234, | Xyloglucan endotransglucosylase/hydrolase proteins |
|  | LOC115714877 |  |
| **Signaling & Defense** | LOC115718923 | Calcium-binding protein CML37 |
| **Signaling & Defense** | LOC115699995 | Probable protein kinase At2g41970 |
| **Signaling & Defense** | LOC115717279,<br>LOC115716798 | Ethylene-responsive transcription factors |
| **Signaling & Defense** | LOC115702722 | WRKY transcription factor 15 |
| **Signaling & Defense** | LOC115700633 | NDR1/HIN1-like protein 3 |
| **Stress & Detoxification** | LOC115712390 | SULFUR DEFICIENCY-INDUCED 1 protein |
| **Stress & Detoxification** | LOC115722134 | Cytochrome P450 94A1-like |
| **Stress & Detoxification** | LOC115721051,<br>LOC115713627 | Zinc finger proteins (ZAT12, ZAT10-like) |
| **Stress & Detoxification** | LOC115720647 | Germin-like protein subfamily 1 member 13 |
| **RNA & Transcriptional Regulation** | LOC115719700 | Probable CCR4-associated factor 1 homolog 9 |
| **RNA & Transcriptional Regulation** | LOC115716702 | DEAD-box ATP-dependent RNA helicase 48 |
| **Metabolism** | LOC115722728 | Sugar transporter ERD6-like 16 |
| **Metabolism** | LOC115702961 | Cysteine proteinase RD21A-like |
| **Metabolism** | LOC115697811 | Allantoinase |

**TABLE 3-B.** Down-regulated Genes in infected HLVd Tissue. Genes significantly downregulated (Greater than -3 log fold change) in infected tissue libraries.

| **Activity** | **Gene ID** | **Annotated Gene Name** |
| --- | --- | --- |
| **Cell Wall & Structural Components** | LOC115695646 | Glycine-rich cell wall structural protein 2 |
| **Protein Regulation & Modification** | LOC115712493 | Probable E3 ubiquitin-protein ligase XERICO |
| **Protein Regulation & Modification** | LOC115717232 | Protein phosphatase 2C 51 |
| **Transcription & Gene Expression** | LOC115711395 | Nuclear transcription factor Y subunit A-1 |
| **Lipid Transfer & Metabolism** | LOC115698126,<br>LOC115724726,<br>LOC115722891 | Non-specific lipid-transfer proteins |
| **Lipid Transfer & Metabolism** | LOC115720958 | (R)-specific enoyl-CoA hydratase |
| **Defense & Stress** | LOC115698859 | Mannose/glucose-specific lectin-like |
| **Defense & Stress** | LOC115705814 | Pathogenesis-related protein 1 |
| **Photosynthesis & Energy** | LOC115716632 | Dihydrodipicolinate reductase-like protein CRR1, chloroplastic |
| **Photosynthesis & Energy** | LOC115725758 | Ferredoxin--NADP reductase, root isozyme, chloroplastic-like |

Nanopore cDNA Sequencing of HLVd positive and negative meristematic tissue 6 Replicate libraries of cDNA were prepared from previously treated, regenerated, and positive and negative testing Anna Lee meristematic plant tissue. Positive and negative plants were also tested prior to sequencing to confirm HLVd status. Anna Lee was chosen because it showed an intermediate capacity for viroid clearance relative to the other 4 cultivars in the study, and we sought to investigate the differential gene activity in positive and negative individuals of this cultivar. The positive and negative cDNA libraries were prepared and each Library barcoded for sequencing on a R.9.4.1 flow cell on MinIon following the cDNA Sequencing protocol (SQK-PCS109) (Oxford Nanopore, UK) using basic PC with the following specifications: Ryzen 7 5800X 8-Core Processor 3.80 GHz, 16.0 GB RAM, 64-bit windows operating system. The data is uploaded to Genbank under Bio project number PRJNA1083500 and SRA reads include 6 replicate clean Anna Lee tissue libraries (SAMN40250909-SAMN40250914) and 6 replicate infected Anna Lee tissue libraries (SAMN40250915-SAMN40250920).

### Basecalling & Trimming

Reads obtained from the MinIon sequencer were basecalled using the high accuracy (HA) model of Guppy (Oxford Nanopore, UK) using the GPU basecalling mode. These reads were trimmed using default parameters to remove the adapter sequences and provide only reads passing the default quality filter of Guppy. They were then demultiplexed using Guppy Barcoder (Oxford Nanopore, UK) and then combined using a python script to have a single set of trimmed reads for each replicate to be input into the differential expression pipeline.

### Differential Expression Analysis

To evaluate the expression differences between infected and non-infected plants, the reads were used to perform a reference guided differential expression analysis using the Epi2Me pipeline (https://labs.epi2me.io/wfindex/). 6 replicates of each condition were mapped to the CS10 (28) RefSeq genome (Accession # GCF_900626175.2) using minimap2 (15). The reads were then quantified using salmon (29), and identified transcripts were normalized across libraries before being input into DEXseq (30) and DeSeq2 (2) pipelines to perform the differential expression. The results were filtered using the following parameters: +/-3-Logfold Change, <0.001 P-Value, and a <0.01 False Discovery Rate.

## Acknowledgments

The authors are thankful for Cecilia Zapata, who helped with tissue culture guidance during the thermotherapy work. Additionally, the authors would like to acknowledge Jonathan Vaught and Front Range Biosciences for enabling and funding this work throughout the years. Finally, the authors would like to thank Alyssa Burkhardt and Tessa Albrecht for their support to ensure the accuracy and reproducibility of the technology.

## Author contributions

AT conceptualized the experiment, wrote the manuscript, carried out data curation, formal analysis, and data visualization. CP performed the formal analysis of DGE, data visualization, reviewed and edited the manuscript. CZ supported supervision, reviewed, and edited the manuscript, funding acquisition, and project administration. CS performed tissue culture methodology and thermotherapy treatment. RG supervised the work and contributed to writing and editing the final manuscript.

## Supporting Information Captions

Figure S.A. Standard Curve of Dual Interrogation Real-time Assay. Plot of Ct data for multiplex RT-qPCR assay measuring HLVd by two separate probes, AMV or BCTV, and the 26S Cannabis gene as a reference control.

Table S.A.T. Table of Ct Data from Standard Curve of Known Input Concentration of RNA from 1ng to 0.00001ng.

Table S.1.T Table of Raw Ct Data from Figure 1. Ct results from 13 HLVd positive and negative cultivars post thermotherapy with replicates and multiple testing time points for negative testing plantlets.

Table S.2.T Table of Ct Means from HLVd positive tests and results from ANOVA

S.3.T Table of Differential Gene Analysis results with LOC ID, Gene Description, logFC (Fold Change), logCPM (Counts per Million), F (score), PValue, FDR (False Discovery Rate)

## Notes

### Competing Interest Statement

The authors have declared no competing interest.

## References

1. Adkar-Purushothama CR, Sano T, Perreault JP. Hop Latent Viroid: A Hidden Threat to the Cannabis Industry. Viruses. 2023;15(3).

2. Love MI, Huber W, Anders S. Moderated estimation of fold change and dispersion for RNA-seq data with DESeq2. Genome Biol. 2014;15(12):550.

3. Warren JG, Mercado J, Grace D. Occurrence of Hop Latent Viroid Causing Disease in Cannabis sativa in California. Plant Disease. 2019;103(10):2699-.

4. Steger G, Riesner D. Viroid research and its significance for RNA technology and basic biochemistry. Nucleic Acids Res. 2018;46(20):10563–76.

5. Puchta H, Ramm K, Sanger HL. The molecular structure of hop latent viroid (HLV), a new viroid occurring worldwide in hops. Nucleic Acids Res. 1988;16(10):4197–216.

6. Punja Z, Kahl D, Reade R, Xiang Y, Munz J, Nachappa P. Challenges to Cannabis sativa Production from Pathogens and Microbes-the Role of Molecular Diagnostics and Bioinformatics.. Int J Mol Sci 2024;25(14).

7. Bektaş A, Hardwick KM, Waterman K, Kristof J. Occurrence of Hop Latent Viroid in Cannabis sativa with Symptoms of Cannabis Stunting Disease in California. Plant Disease. 2019;103(10).

8. Chiginsky J, Langemeier K, MacWilliams J, Albrecht T, Cranshaw W, Fulladolsa AC, et al. First Insights Into the Virus and Viroid Communities in Hemp (Cannabis sativa). Frontiers in Agronomy. 2021;3.

9. Jarugula S, Wagstaff C, Mitra A, Crowder D, Gang D, Rayapati N. First reports of Beet curly top virus, Citrus yellow vein-associated virus, and Hop latent viroid in industrial hemp (Cannabis sativa) in Washington State. Plant Dis. 2023.

10. Schoener JL. Molecular Identification of Plant Pathogens Affecting Cannabis sativa Crops in Nevada: University of Nevada, Reno; 2021.

11. Punja ZK. Emerging diseases of Cannabis sativa and sustainable management. Pest Manag Sci. 2021;77(9):3857–70.

12. Fernandez IMA, Parungao M, Hollin J, Selimotic B, Farrar G, Seyler T, et al. A Novel, Precise and High-Throughput Technology for Viroid Detection in Cannabis (MFDetect(TM)). Viruses. 2023;15(7).

13. Pokorn T, Radisek S, Javornik B, Stajner N, Jakse J. Development of hop transcriptome to support research into host-viroid interactions. PLoS One. 2017;12(9):e0184528.

14. Laimer M, Barba M. Elimination of systemic pathogens by thermotherapy, tissue culture, or in vitro micrografting. Virus and virus-like diseases of pome and stone fruits. 2011:389–93.

15. Li H. Minimap2: pairwise alignment for nucleotide sequences. Bioinformatics. 2018;34(18):3094–100.

16. Ramesh SV, Yogindran S, Gnanasekaran P, Chakraborty S, Winter S, Pappu HR. Virus and Viroid-Derived Small RNAs as Modulators of Host Gene Expression: Molecular Insights Into Pathogenesis. Front Microbiol. 2020;11:614231.

17. Miotti N, Passera A, Ratti C, Dall’Ara M, Casati P. A Guide to Cannabis Virology: From the Virome Investigation to the Development of Viral Biotechnological Tools. Viruses. 2023;15(7).

18. Sirangelo TM, Ludlow RA, Spadafora ND. Molecular Mechanisms Underlying Potential Pathogen Resistance in Cannabis sativa. Plants (Basel). 2023;12(15).

19. Kappagantu M, Nelson ME, Bullock JM, Kenny ST, Eastwell KC. Hop stunt viroid: Effects on vegetative growth and yield of hop cultivars, and its distribution in central Washington state. Plant disease. 2017;101(4):607–12.

20. Matousek J, Trnena L, Svoboda P, Oriniakova P, Lichtenstein CP. The gradual reduction of viroid levels in hop mericlones following heat therapy: a possible role for a nuclease degrading dsRNA. Biol Chem Hoppe Seyler. 1995;376(12):715–21.

21. Matoušek J. Hop latent viroid (HLVd) microevolution: an experimental transmission of HLVd “thermomutants” to solanaceous species. Biologia plantarum. 2003;46:607–10.

22. Narváez-Barragán DA, Tovar-Herrera OE, Guevara-García A, Serrano M, Martinez-Anaya C. Mechanisms of plant cell wall surveillance in response to pathogens, cell wall-derived ligands and the effect of expansins to infection resistance or susceptibility. Frontiers in Plant Science. 2022;13:969343.

23. Khuu NL. Titer and distribution of hop stunt viroid and hop latent viroid infecting hops: Washington State University; 2021.

24. Itaya A, Zhong X, Bundschuh R, Qi Y, Wang Y, Takeda R, et al. A structured viroid RNA serves as a substrate for dicer-like cleavage to produce biologically active small RNAs but is resistant to RNA-induced silencing complex-mediated degradation. J Virol. 2007;81(6):2980–94.

25. Bhojwani SS, Razdan MK. Plant tissue culture: theory and practice: Elsevier; 1986.

26. Leifert C, Ritchie J, Waites W. Contaminants of plant-tissue and cell cultures. World Journal of Microbiology and Biotechnology. 1991;7(4):452–69.

27. Ahmed M, Kim DR. pcr: an R package for quality assessment, analysis and testing of qPCR data. PeerJ. 2018;6:e4473.

28. Grassa CJ, Weiblen GD, Wenger JP, Dabney C, Poplawski SG, Timothy Motley S, et al. A new Cannabis genome assembly associates elevated cannabidiol (CBD) with hemp introgressed into marijuana. New Phytologist. 2021;230(4):1665–79.

29. Patro R, Duggal G, Love MI, Irizarry RA, Kingsford C. Salmon provides fast and bias-aware quantification of transcript expression. Nat Methods. 2017;14(4):417–9.

30. Anders S, Reyes A, Huber W. Detecting differential usage of exons from RNA-seq data. Genome Res. 2012;22(10):2008–17.

