## Supplemental for "Differential Expression Analysis of Cannabis sativa response to thermotherapy of Hops Latent Viroid (HLVd) infection and clearance of the viroid in Tissue Culture Micropropagation"

RT-qPCR Multiplex Detection (Supplemental)

As described in US patent # US11739374B2, 1µL of normalized total RNA (normalized to 1ng/µL) was tested with HLVd/26S and HLVd/BCTV/26S RT-qPCR multiplex assays. A HLVd/26S multiplex assay was prepared by formulating QuantNova one step RT-qPCR mastermix (Qiagen, Germany) with the B-F HLVd primer pair with the HLVd probe p4 labeled with 6-FAM in multiplex and HLVd probe p2 labeled with ROX NHS and 1 internal positive control 26S ribosomal RNA primer pair with 26S probe p1 labeled with SUN. A second HLVd/BCTV/26S multiplex was prepared by formulating QuantNova one step RT-qPCR mastermix with the B-F HLVd primer pair with the HLVd probe p4 labeled with 6-FAM in multiplex and HLVd probe p2 labeled with ROX NHS, the BCTV DRP MP primer pair with BCTV DRP MP Probe 2 labeled with Cy5 and BCTV DRP MP Probe 1 labeled with TAMRA NHS, and 1 internal positive control 26S ribosomal RNA primer pair with 26S probe p1 labeled with SUN.

RT-qPCR assay validation (Supplemental)

To demonstrate robust qualitative sensitivity and specificity for detection of HLVd, BCTV, and AMV in a RT-qPCR assay, both positive and negative Cannabis RNA pools, an RNA standard curve, and a no template control were used as input and assayed with technical replicates in the form of duplicates. In the HLVd AMV 5 Target Multiplex both technical replicates for the 1ng Cannabis RNA pool that was HLVd and AMV Negative, observed a signal for the internal positive control 26S with a Ct value crossing threshold indicating a successful RT-qPCR reaction and no Ct values for HLVd or AMV were observed. In both technical replicates of the 1ng Cannabis RNA pool that was HLVd and AMV positive, a signal for the internal positive control 26S was observed with a Ct value crossing threshold indicating a successful RT-qPCR reaction as well as Ct values for HLVd p2 and p4 and AMV A and AMV B were observed indicating dual interrogation positive detection for HLVd and AMV in the positive pool. In the standard curve reaction, dual positive signals were observed for HLVd p4 and p2 down to 100fg and AMV A Sensitivity down to 10fg and AMV B sensitivity down to 100fg and 26S sensitivity down to 10fg. No Ct value observations were observed for any probe in the no template control. Results illustrated in slide 1-2 “20210128 HLVd AMV 26S and HLVd BCTV and 26S 5Plex Plots”. (Figure S.A ) In the HLVd BCTV 5 Target Multiplex, both technical replicates for the 1ng Cannabis RNA pool that was HLVd and BCTV Negative, observed a signal for the internal positive control 26S with a Ct value crossing threshold indicating a successful RT-qPCR reaction and no Ct values for HLVd or BCTV were observed. In both technical replicates of the 1ng Cannabis RNA pool that were HLVd and BCTV positive, a signal for the internal positive control 26S was observed with a Ct value crossing threshold indicating a successful RT-qPCR reaction as well as Ct values for HLVd p2 and p4 and BCTV DRP MP Probe 1 and 2 were observed indicating dual interrogation positive detection for HLVd and BCTV in the positive pool. In the standard curve reaction, dual positive signals were observed for HLVd p4 and p2 down to 100fg and BCTV DRP MP Probe 1 and 2 Sensitivity down 1pg and 26S sensitivity down to 10fg. No Ct value observations were observed for any probe in the no template control. Results illustrated in slide 1-2 “20210128 HLVd AMV 26S and HLVd BCTV and 26S 5Plex Plots”. (Figure S.A )

Table S.2.T Statistics

| MeanCt | Cultivar | HLVd | Reference |  |
| --- | --- | --- | --- | --- |
|  | Abigail | 23.35671 | 21.84543 |  |
|  | Amber | 27.19971 | 15.61351 |  |
|  | AnnaLee | 28.02563 | 22.01027 |  |
|  | Athena | 19.98396 | 21.12142 |  |
|  | EarlyPearly | 31.87651 | 16.48788 |  |
|  | FRB1.4 | 32.09273 | 17.01117 |  |
|  | Hybrid5 | 30.91255 | 18.68844 |  |
|  | Hybrid9 | 25.79425 | 15.84535 |  |
|  | S1.4 | 31.12595 | 16.83204 |  |
|  | Wife | 21.4862 | 21.15476 |  |
|  | PureCBG | 31.35479 | 14.76573 |  |
| Analysis of Variance Table | RiverRock | df | F-value | pvalue |
|  | Cultivar | 11 | 127.8 | 0.001 |

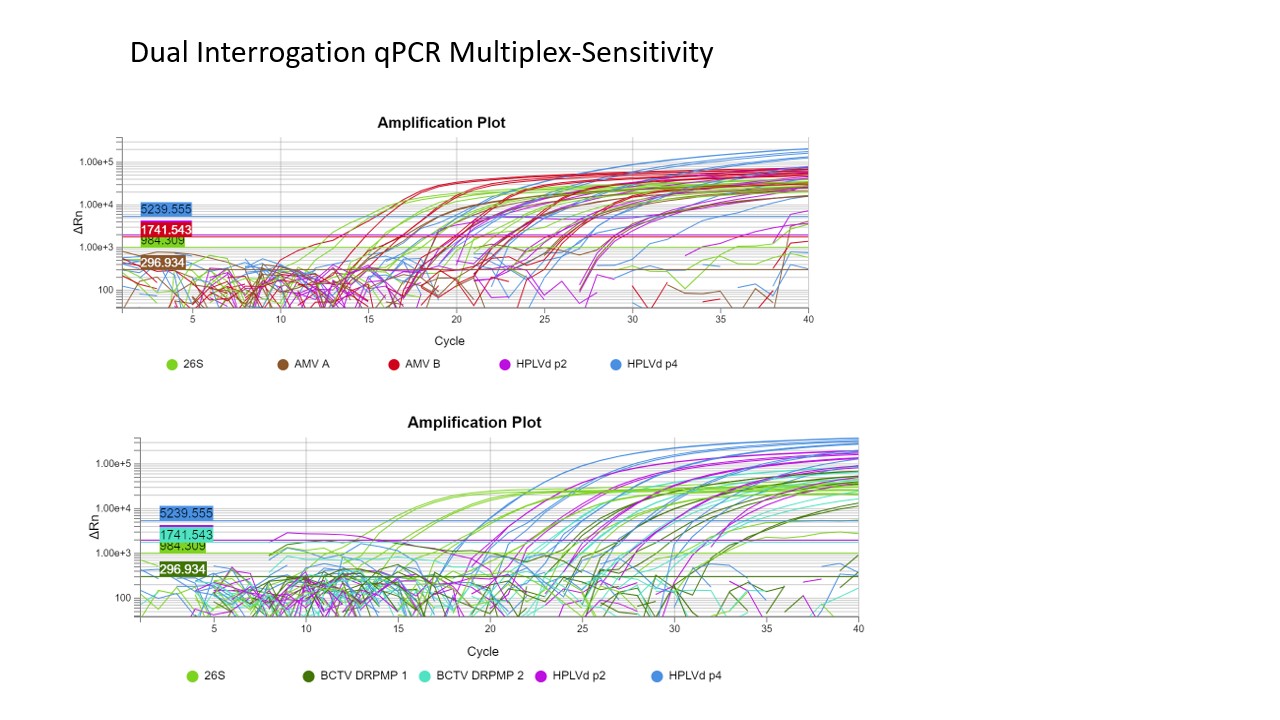

Figre S.A

Supplementary Material

Table S.3.T

| **LOC#** | **Gene Description** | **logFC** | **logCPM** | **F** | **PValue** | **FDR** |
| --- | --- | --- | --- | --- | --- | --- |
| LOC115705195 | protein ALP1-like | 6.430861423 | 5.344049714 | 68.23018389 | 1.46E-16 | 2.26E-14 |
| LOC115700823 | uncharacterized LOC115700823 | 6.33083608 | 5.268071893 | 62.91081655 | 2.18E-15 | 2.79E-13 |
| LOC115724771 | uncharacterized LOC115724771 | 6.172622734 | 5.174321985 | 56.40544158 | 5.92E-14 | 6.49E-12 |
| LOC115716935 | expansin-like B1 | 5.681859063 | 4.864839425 | 38.84583823 | 4.60E-10 | 3.14E-08 |
| LOC115718923 | calcium-binding protein CML37 | 5.186750092 | 4.580684157 | 26.63530977 | 2.46E-07 | 1.05E-05 |
| LOC115699995 | probable protein kinase At2g41970 | 5.170891388 | 4.578503137 | 26.4320226 | 2.73E-07 | 1.16E-05 |
| LOC115722728 | sugar transporter ERD6-like 16 | 4.825395667 | 5.456699778 | 61.84260459 | 3.74E-15 | 4.73E-13 |
| LOC115719700 | probable CCR4-associated factor 1 homolog 9 | 4.817066862 | 8.448739777 | 648.476474 | 8.76E-143 | 3.82E-139 |
| LOC115712390 | protein SULFUR DEFICIENCY-INDUCED 1 | 4.807715476 | 4.409006143 | 20.4122923 | 6.25E-06 | 0.000206584 |
| LOC115696797 | F-box protein At1g22220 | 4.69961866 | 5.378054075 | 55.80789879 | 8.03E-14 | 8.64E-12 |
| LOC115722134 | cytochrome P450 94A1-like | 4.653890585 | 4.337301923 | 18.3501562 | 1.84E-05 | 0.00054881 |
| LOC115723779 | uncharacterized LOC115723779 | 4.449444822 | 4.259327454 | 16.0608963 | 6.14E-05 | 0.001589171 |
| LOC115721452 | protein EXORDIUM-like 1 | 4.44862595 | 7.481373336 | 309.0369013 | 4.06E-69 | 5.06E-66 |
| LOC115704140 | strigolactone esterase D14-like | 4.32937486 | 4.218289105 | 14.8801501 | 0.000114603 | 0.002781765 |
| LOC115720647 | germin-like protein subfamily 1 member 13 | 4.225796688 | 4.178214888 | 13.97956842 | 0.000184867 | 0.004195135 |
| LOC115716702 | probable DEAD-box ATP-dependent RNA helicase 48 | 4.20315134 | 4.176647184 | 13.83892265 | 0.00019923 | 0.004474455 |
| LOC115724651 | probable xyloglucan endotransglucosylase/hydrolase protein 23 | 4.138802439 | 6.391462765 | 124.1451791 | 8.01E-29 | 2.45E-26 |
| LOC115704623 | uncharacterized LOC115704623 | 4.105124763 | 4.136071573 | 12.9892576 | 0.00031337 | 0.006417649 |
| LOC115702961 | cysteine proteinase RD21A-like | 4.099396202 | 4.135946261 | 12.96305495 | 0.000317786 | 0.006477641 |
| LOC115725776 | zinc finger Ran-binding domain-containing protein 2 | 4.075431279 | 4.13432718 | 12.82189889 | 0.000342679 | 0.006920288 |
| LOC115721051 | zinc finger protein ZAT12 | 4.056959384 | 5.257784559 | 42.74995228 | 6.24E-11 | 4.83E-09 |
| LOC115725683 | probable amino acid permease 7 | 4.048590264 | 5.241092871 | 42.08907856 | 8.74E-11 | 6.54E-09 |
| LOC115697811 | allantoinase | 3.820885587 | 6.002639743 | 81.13849244 | 2.12E-19 | 4.11E-17 |
| LOC115711935 | uncharacterized LOC115711935 | 3.820198334 | 4.482782627 | 16.88920931 | 3.96E-05 | 0.001084648 |
| LOC115724781 | rRNA 2'-O-methyltransferase fibrillarin-like | 3.692470013 | 5.015560793 | 30.28358153 | 3.74E-08 | 1.84E-06 |
| LOC115700633 | NDR1/HIN1-like protein 3 | 3.620991722 | 6.671812509 | 134.1030346 | 5.32E-31 | 1.69E-28 |
| LOC115717279 | ethylene-responsive transcription factor 5 | 3.60945073 | 5.846845832 | 65.90483426 | 4.76E-16 | 6.86E-14 |
| LOC115706272 | uncharacterized LOC115706272 | 3.527960582 | 4.372427558 | 13.21780802 | 0.000277382 | 0.005831369 |
| LOC115702722 | probable WRKY transcription factor 15 | 3.47945479 | 8.369514156 | 431.5891074 | 9.58E-96 | 2.39E-92 |
| LOC115695862 | pentatricopeptide repeat-containing protein At5g11310, mitochondrial | 3.305590716 | 4.830730495 | 21.06010955 | 4.45E-06 | 0.000153714 |
| LOC115713627 | zinc finger protein ZAT10-like | 3.303749147 | 6.970127906 | 151.4219512 | 8.76E-35 | 3.32E-32 |
| LOC115716798 | ethylene-responsive transcription factor 5 | 3.244104152 | 7.395962032 | 203.2576784 | 4.31E-46 | 2.95E-43 |
| LOC115721738 | protein C2-DOMAIN ABA-RELATED 4 | 3.196421283 | 5.128797988 | 28.32815388 | 1.03E-07 | 4.74E-06 |
| LOC115712234 | probable xyloglucan endotransglucosylase/hydrolase protein 6 | 3.188792517 | 11.31711226 | 1318.349108 | 1.40E-287 | 2.43E-283 |
| LOC115714877 | probable xyloglucan endotransglucosylase/hydrolase protein 33 | 3.128105364 | 6.416584185 | 90.32802064 | 2.04E-21 | 4.62E-19 |
| LOC115697909 | kiwellin-like | 3.119025383 | 4.481861296 | 12.88221417 | 0.00033181 | 0.006724178 |
| LOC115710253 | mitogen-activated protein kinase kinase 9-like | 3.075472083 | 7.351016894 | 182.1349955 | 1.74E-41 | 9.26E-39 |
| LOC115717356 | heavy metal-associated isoprenylated plant protein 39 | 3.001107161 | 7.118165293 | 149.0360079 | 2.91E-34 | 1.08E-31 |
| LOC115705702 | uncharacterized LOC115705702 | -3.008935039 | 4.807409744 | 13.65299624 | 0.000219957 | 0.004809808 |
| LOC115695646 | glycine-rich cell wall structural protein 2 | -3.012707699 | 5.716851784 | 35.47844898 | 2.58E-09 | 1.52E-07 |
| LOC115700980 | putative methyltransferase DDB_G0268948 | -3.054745085 | 5.554930766 | 30.99801417 | 2.59E-08 | 1.30E-06 |
| LOC115717038 | uncharacterized LOC115717038 | -3.081851948 | 6.708467335 | 89.38018295 | 3.30E-21 | 7.36E-19 |
| LOC115698859 | mannose/glucose-specific lectin-like | -3.141541858 | 6.563420756 | 81.00879739 | 2.27E-19 | 4.35E-17 |
| LOC115724954 | uncharacterized LOC115724954 | -3.181433281 | 4.892596418 | 16.21024352 | 5.67E-05 | 0.001486247 |
| LOC115712493 | probable E3 ubiquitin-protein ligase XERICO | -3.356971686 | 6.530671713 | 84.6908396 | 3.53E-20 | 7.14E-18 |
| LOC115711395 | nuclear transcription factor Y subunit A-1 | -3.411189807 | 5.040627554 | 20.68574162 | 5.42E-06 | 0.000183185 |
| LOC115716632 | dihydrodipicolinate reductase-like protein CRR1, chloroplastic | -3.41557407 | 5.028297388 | 20.57204667 | 5.75E-06 | 0.000192602 |
| LOC115704444 | stemmadenine O-acetyltransferase | -3.421741826 | 4.637250175 | 13.02631469 | 0.00030723 | 0.006306726 |
| LOC115701693 | uncharacterized LOC115701693 | -3.426656179 | 4.62002809 | 12.90541918 | 0.000327722 | 0.006656811 |
| LOC115710355 | uncharacterized LOC115710355 | -3.432699486 | 4.641908222 | 13.13467507 | 0.000289963 | 0.006030399 |
| LOC115699374 | putative uncharacterized protein DDB_G0282499 | -3.439810385 | 4.642841646 | 13.19996391 | 0.000280035 | 0.005865922 |
| LOC115698126 | non-specific lipid-transfer protein 1-like | -3.551050758 | 5.921125772 | 51.74679964 | 6.34E-13 | 6.39E-11 |
| LOC115724726 | non-specific lipid-transfer protein 8 | -3.563811513 | 4.702790157 | 14.74369226 | 0.000123202 | 0.002957536 |
| LOC115704261 | uncharacterized LOC115704261 | -3.690418432 | 4.77800976 | 16.65500946 | 4.49E-05 | 0.001213782 |
| LOC115717232 | protein phosphatase 2C 51 | -3.703855671 | 4.7781708 | 16.79445077 | 4.17E-05 | 0.001134829 |
| LOC115700445 | major latex allergen Hev b 5 | -3.773694696 | 6.287954105 | 77.25149721 | 1.52E-18 | 2.70E-16 |
| LOC115719589 | uncharacterized LOC115719589 | -3.869585948 | 5.309219289 | 31.49610913 | 2.00E-08 | 1.03E-06 |
| LOC115722891 | non-specific lipid-transfer protein 4.1 | -3.92259793 | 4.889103244 | 20.37469967 | 6.37E-06 | 0.000209886 |
| LOC115705814 | pathogenesis-related protein 1 | -4.258601993 | 5.095234206 | 27.59960992 | 1.49E-07 | 6.69E-06 |
| LOC115703097 | glutaredoxin-C13 | -4.260750032 | 4.353168273 | 12.27451871 | 0.0004593 | 0.008796354 |
| LOC115712485 | uncharacterized LOC115712485 | -4.347316447 | 4.388739567 | 12.9900545 | 0.000313237 | 0.006417649 |
| LOC115725758 | ferredoxin--NADP reductase, root isozyme, chloroplastic-like | -4.35705942 | 4.389864966 | 13.0535332 | 0.000302798 | 0.00623778 |
| LOC115707893 | uncharacterized LOC115707893 | -4.361466698 | 4.39058831 | 13.0854909 | 0.000297676 | 0.006145344 |
| LOC115708714 | uncharacterized LOC115708714 | -4.423321694 | 4.422675306 | 13.72093313 | 0.000212143 | 0.004680034 |
| LOC115700634 | uncharacterized LOC115700634 | -4.45355177 | 4.427064563 | 13.94959092 | 0.000187838 | 0.004245974 |
| LOC115699918 | uncharacterized LOC115699918 | -4.508132649 | 4.457545671 | 14.56219163 | 0.000135655 | 0.003207867 |
| LOC115707556 | probable inositol oxygenase | -4.564396798 | 7.273835543 | 209.6179591 | 1.77E-47 | 1.34E-44 |
| LOC115697023 | putative lipid-transfer protein DIR1 | -4.567416777 | 6.183936621 | 83.61759048 | 6.07E-20 | 1.22E-17 |
| LOC115720958 | (R)-specific enoyl-CoA hydratase | -4.681309655 | 4.528054322 | 16.36771889 | 5.22E-05 | 0.001386499 |
| LOC115702728 | probable LRR receptor-like serine/threonine-protein kinase At4g37250 | -4.698884294 | 4.551271971 | 16.69811991 | 4.38E-05 | 0.001190207 |
| LOC115704859 | S-norcoclaurine synthase | -4.736341956 | 4.57765669 | 17.22976711 | 3.31E-05 | 0.000921052 |
| LOC115715608 | granule-bound starch synthase 1, chloroplastic/amyloplastic | -4.915495941 | 4.666344558 | 19.60489192 | 9.53E-06 | 0.000300695 |
| LOC115709904 | phosphatidylinositol 4-phosphate 5-kinase 9 | -5.116582588 | 4.75929397 | 22.53750011 | 2.06E-06 | 7.62E-05 |
| LOC115721929 | protein RSI-1 | -5.35211935 | 4.904791868 | 27.17225581 | 1.86E-07 | 8.20E-06 |
